# The Role of Glycocalyx Steric Effects on Viral Endocytosis*

**DOI:** 10.1101/2022.02.12.480189

**Authors:** Joseph Buckley, Giuseppe Battaglia

**Affiliations:** Department of Chemistry and Institute for the Physics of Living Systems, University College London, London, UK. Institute for Bioengineering of Catalonia (IBEC), The Barcelona Institute for Science and Technology (BIST), Barcelona, Spain

## Abstract

Understanding the mechanisms of viral entry is key to aiding the development of effective anti-viral treatments. In this work, we produce a simple thermodynamic model of viral entry, which is able to predict the differences between viruses. We also show that the glycocalyx, which is often neglected in studies of cell-entry, plays a key role, and the density and structure of the glycocalyx can determine whether or not a virus can enter the cell. We further find that co-receptors play not only a signalling role, but an important thermodynamic role in viral entry. We then show that this model can be used to calculate the cell-entry probabilities of a given virus, which can shine a light on the clinical observations associated with a virus.

## INTRODUCTION

The structure of viruses includes genetic material, either DNA or RNA, surrounded by a protein shell or capsid. Enveloped viruses are also covered by a lipid membrane derived from the host cell.

Although made of the same components as living cells, viruses are incapable of self-replicating and thus ‘infect’ cells hijacking their synthetic machinery in order to reproduce. The proteins that coat viruses or spikes, often stabilised by complex glycans, have evolved to bind to several receptors expressed on the host cells

Viruses can form multiple bonds with the host cell, a process called multivalent binding. Once attached to the cell membrane, a virus may enter the cell.

Viruses infect the cell through two different pathways: endocytosis, where the virus is wrapped by the cell and internalised, and membrane fusion, where the viral membrane fuses with the cell membrane.

During endocytosis, the host cells membrane deform to wrap the virus fully. Such a process is driven by the formation of additional bonds to the virus. As the membrane deforms, the strain increases on the junction between the deformed and non-deformed part of the membrane; eventually, this strain becomes so great it causes the membrane to pinch off in a process aided by specialised proteins, called fission. The membrane then returns to its original state without losing integrity along the process, with the virus internalised inside a trafficking vesicle. Frequently, internal cell mechanisms help along this process. For example, among all the different endocytic proteins, those that comprise Bin/amphiphysin/Rvs (BAR) domains have an anisotropic shape that can bind to the inside of the curved membrane [1], which may make the deformed state more energetically favourable. Such a step is a key part of clathrin-mediated endocytosis, the most studied form of endocytosis [2, 3].

In addition to this, most cells are coated by a dense network of highly charged glycosaminoglycans (GAGs) chains, with the most abundant being the Heparan Sulfate (HS). These polymers are tethered to the cell membrane via proteins, appropriately named proteoglycans. The resulting chemical cushion as thick as hundreds of nanometers acts as the first defence for cells, and it is collectively known as the glycocalyx. Viruses and other pathogens will need to overcome such a barrier to access the cell interior during infection.

However, due to the large size of these sugars, they can play a key role in cell-virus interaction due to the steric repulsion they induce upon binding[4]. In most studies on endocytosis, the role of GAGs such as HS is neglected. While, for certain viruses, this is likely not to make a large impact, it has been shown that HS plays a key role in the attachment of both SARS-CoV-2[5], and HIV-1 [6, 7]. Therefore, understanding the steric effects arising from HS chains is critical to understanding the endocytosis of viruses. Additionally, a HS analogue, heparin, can inhibit multiple viruses, including SARS-CoV-2[8] and HIV-1[9]. Therefore, a complete understanding of the role of HS in viral entry may explain why heparin is an effective treatment. In this work, we demonstrate that the steric repulsion arising from GAGs, and HS in particular, favours membrane deformation and thus endocytosis.

## THEORY

The total energy of interaction between a virus and a cell can be decomposed into an attractive binding component, *E_att_*, four repulsive components: the membrane bending energy *E_b_*, the membrane tension energy, *E_T_*, and an energy describing the steric repulsion due to the presence of HS, *E_rep_*. This gives

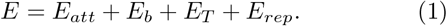

All contributions to the energy can be expressed as a function of engulfment depth, *h*, the maximum distance of membrane perturbation. This model will assume a single bond between the surface and virus has already been formed. For more detail on the attachment of viruses, see our previous work[4].

### Membrane Bending Energy

The membrane bending energy depends on the bending modulus *κ*, the virion radius *R*, and the spontaneous bending *C*, which accounts for deformation induced by intracellular mechanisms such as BAR protein aggregation or the formation of clathrin-coated pits. We can express the spontaneous bending as some fraction of the virion radius, 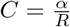, to give

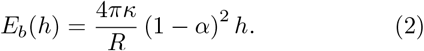

The spontaneous bending is often taken to be zero implying that intercellular processes are not necessary for endocytosis. There is very little evidence suggesting such a mechanism, which would need to be driven either by long-range attractive interactions or by random fluctuations in the membrane. These interactions/fluctuations would have to bridge a distance larger than twenty nanometres, which is highly unlikely. Therefore, we cannot neglect spontaneous bending.

### Membrane Tension Energy

The membrane tension energy represents the membrane resistance to external forces. It is dependent on the membrane tension, *σ* (typically around 0.005 kT nm^−2^), and is given by [10, 11]

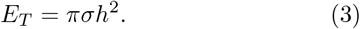

### Attractive Energy

In general, calculation of the attractive energy for multivalent binding requires consideration of the superposition of possible states[12]. However, as viruses typically form very strong bonds to their targetted receptors, the binding energy can be approximated in terms of the receptor-spike binding energy in solution Δ*G*, and the number of bonds λ, as

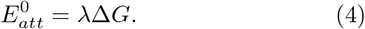

To account for the non-uniform distribution of both receptors and spikes, we perform a Poisson averaging to give

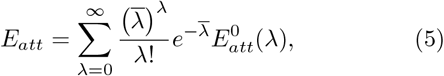

where 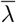 is the average number of bonds formed.

Due to the large binding energy, the average number of bonds formed will roughly be the lowest of either the number of spikes or the number of receptors. For most viruses, this will be the number of receptors. However, for viruses with a very low number of spikes, for example, HIV-1, the number of bonds is limited by the number of spikes.

The average number of spikes and receptors, 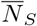 and 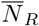, are dependent on the virus-membrane contact area *A* = 2*πhR*, and can be expressed in terms of their respective densities *ρ*_(*S/R*)_,

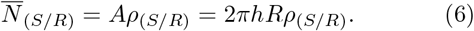

### Repulsive Energy

The repulsive energy that acts on the virus arises from interaction with long-chain sugars that coat the cell surface. In order to bind to the surface, the virus must insert itself into this brush. For a flat surface, this energy can be modelled in terms of the volume of the inserted virus portion, *V*, the insertion parameter, *δ*, and the density of chains, *ρ_HS_*: [13]

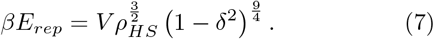

The insertion parameter quantifies how far into the brush the virus has inserted, and can be defined in terms of the lengths of the GAGs — *d_GAG_*, the spikes — *d_S_*, and the receptors — *d_R_*:

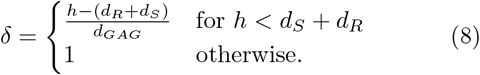

The inserted volume portion can be calculated as,

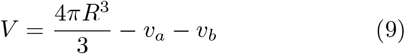

where *v_a_* represents any part of the sphere above the brush, and is non-zero if *d_a_* = 2*R* + *d_R_* + *d_S_* – *h* – *d_GAG_* is greater than zero, such that

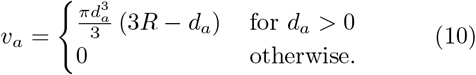

Likewise, *v_b_* represents the any part of the sphere below the brush, and is defined as

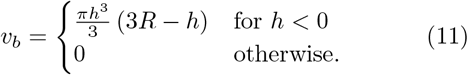

## RESULTS

In this section, we will focus on the two viruses currently causing pandemics — HIV-1 and SARS-CoV-2. These viruses share a lot of similarities, for instance their ability to bind to HS[5, 14]. While both these viruses are often thought to entering the cell by fusion, there is evidence that both can enter via endocytosis[15], with HIV-1 using endocytosis in the absence of the CD4 receptor [3, 16], with some work arguing that HIV-1’s main entry pathway is through endocytosis[17, 18], however this remains controversial.

The two viruses have very different clinical outcomes: SARS-CoV-2 is a highly transmissible disease, with a typical onset of 5 days, which usually lasts around two weeks with a fatality rate of about 4% [19]; HIV-1, however, has very low transmissibility, with the onset of early symptoms after around 2-4 weeks, followed by a latent period which can last up to 10 years. If untreated, the fatality rate of HIV-1 is around 100%.

### The Role Of HS

HIV-1 has an average radius of 63.25 nm [20], and an average number of spikes of 10 per virion [20, 21]. The spike, which is made up of two proteins, gp120, and gp41, bind primarily to the CD4 receptor, with binding energy estimated to be –19 kT [22]. Upon binding, we also expect the spike to interact with a coreceptor, either CCR5 or CXCR4, with energy of roughly – 17 kT [23]. The typical density of CD4 and the coreceptors are much greater than the density of spikes[24], meaning the number of spikes is the limiting factor in the number of bonds formed.

Likewise, SARS-CoV-2 has an average radius of 45 nm [25], and an average number of spikes of 23 per virion. The spike binds to ACE2 with an affinity of –18 kT and to the co-receptor TMPRSS2 with an affinity of – 20*kT* [4]. We will assume ACE2 is the limiting density, and will take the density to be around 56 *μ*m^−2^ [4].

We will assume both spikes and receptors to have a length of 10 nm. We will assume the HS chains have an average length of 50 nm, and a density of 3000 chains *μ*m^−2^ for the immune cells HIV-1 targets and 5000 chains *μ*m^−2^ for the epithelial cells SARS-CoV-2 targets, which correspond to between 48 and 41 chains per binding site respectively. We will take the membrane tension to be 0.005 nm^−2^ [10]. Finally, we will take the bending modulus to be 18 kT and assume a spontaneous curvature fraction of 1 for HIV-1, and 0.1 for SARS-CoV-2, which reflects the weaker affinity of HIV-1 needs more help from intracellular processes to enter the cell.

Using the model and parameters presented above, we then investigated how the energy of the system changes as a function of the wrapping fraction, 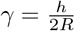 — for both HIV-1 and SARS-CoV-2 in the presence and absence of HS, as can be seen in FIG.2. We see that with a lack of HS chains, an energy well forms, due to the nonlinearity of the membrane tension. For HIV-1 in the absence of HS, this well depth is roughly 28 kT, which could prove an insurmountable barrier to cell entry. However, the presence of HS massively reduces this well to around 8 kT. For SARS-CoV-2, the well has a depth of 12 kT in the absence of HS, which then reduces to 3 kT in the presence of HS.

**FIG. 1.**
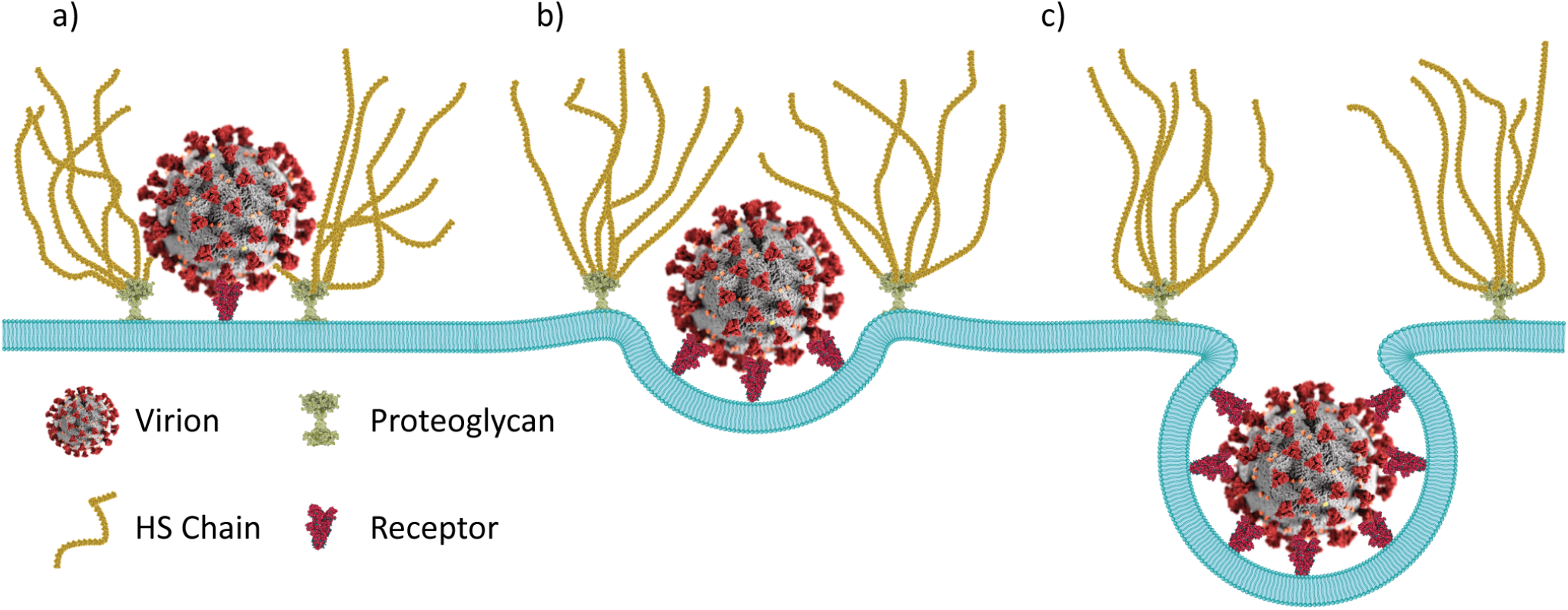
Endocytosis of a virion. **a)** In the first stage of endocytosis, the virion enters the HS brush and binds to the cell surface. **b)** The virion begins to be wrapped by the membrane, and the brush exerts a downward force on the virion. **c)** The virion becomes fully wrapped, and the brush is back in its original state.

**FIG. 2.**
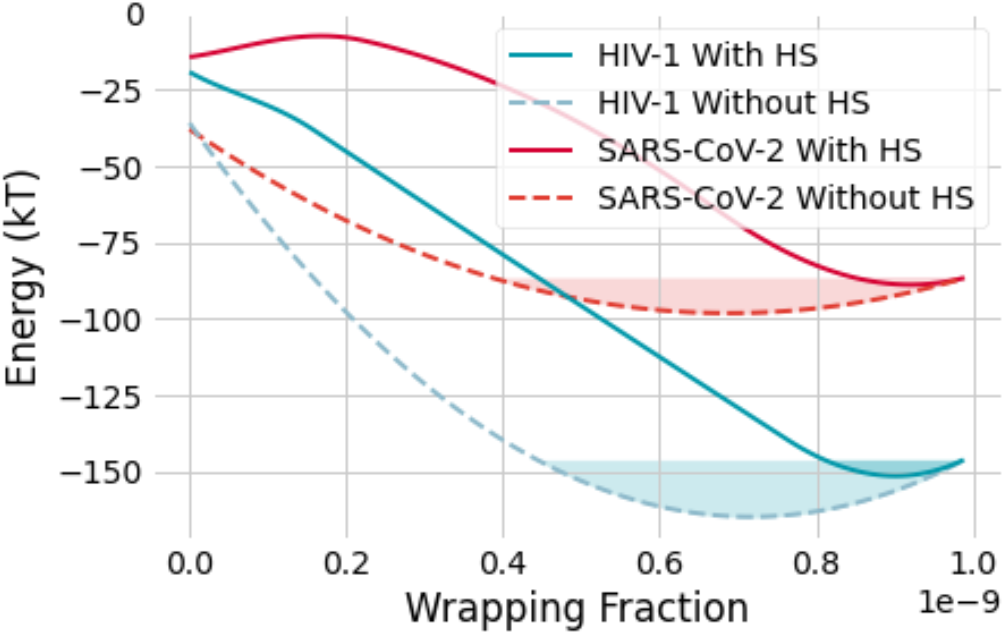
Plots of the energy as a function of wrapping fraction for both HIV-1 and SARS-CoV-2, both in the presence and absence of HS. We can see that in the absence of HS, there is a large potential well, which will provide an energy barrier that the virus will need to overcome in order to endocytose. The presence of HS lowers this barrier.

Such reasoning shows that HS can provide a driving force around endocytosis, allowing a virus to overcome an energy barrier that it previously would be unable to. In FIG.1, we draw a schematic of this process. We can see how the virus disrupts the HS chains upon initial binding. Then as the virus begins to be engulfed, the chains can express a downwards force on the virus until the virus is almost fully wrapped, at which point the chains are no longer disrupted and are back in their original state.

We will define the energy barrier, as the depth of this potential well, as Δ*E* = *E*(*h* = 2*R*) – min(*E*(*h*)), in the presence of the potential well, and Δ*E* = 0 otherwise. In FIG.3b we examine the effect of varying the density of HS on the energy barrier. Below roughly 100 *μ*m^−2^, we see very little effect, as would be expected, as this corresponds to less than one chain per site. However, we see that this energy barrier decreases as the energy increases beyond that range.

**FIG. 3.**
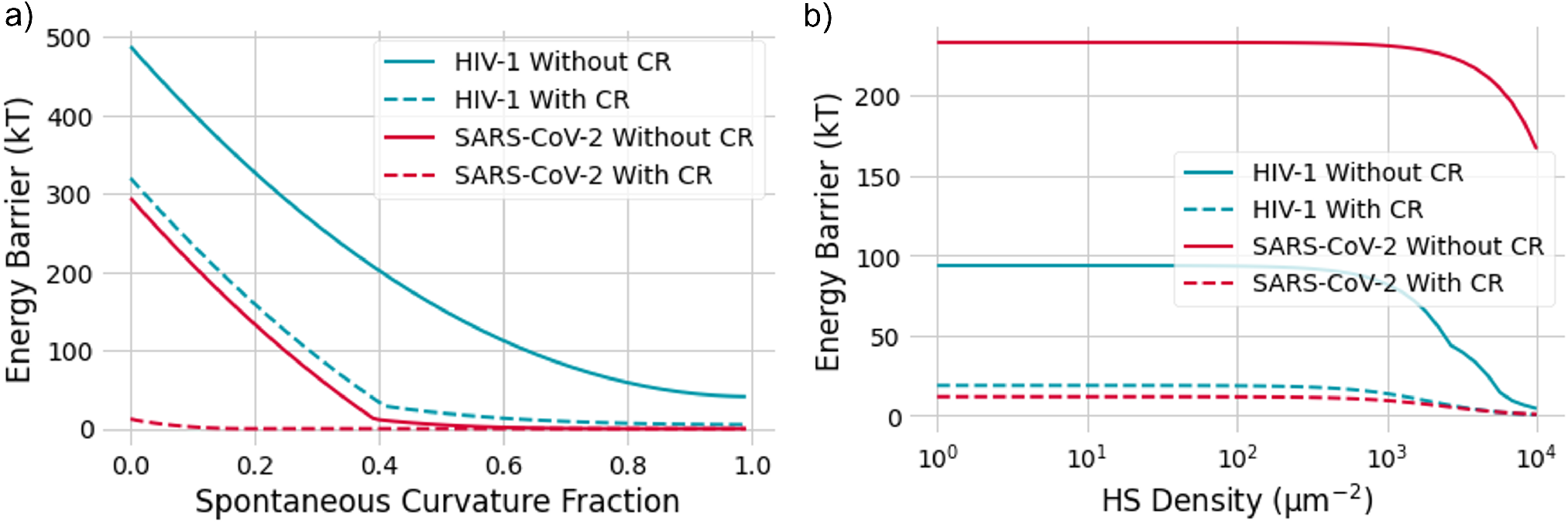
Maximum energy barrier for wrapping as a function of **a)** the spontaneous curvature fracture, *α*, and **b)** the HS density. The solid lines indicate the absence of co-receptors, while the dashed lines indicate the presence of co-receptors. We can observe that in the absence of co-receptors, the energy barrier is often too large for both HIV-1 and SARS-CoV-2, and the presence of co-receptors dramatically decreases this barrier. We see that, in the presence of co-receptors, for HS densities around 500 *μ*m^−2^ the energy barrier begins to drop, this corresponds to roughly 1 chain per virus.

In FIG.3a, we see the effect of varying the spontaneous curvature fraction on the energy barrier. For HIV-1, we observe this energy barrier is huge, around 350 kT, with no spontaneous curvature, while for SARS-CoV-2, we see a significantly lower but still present barrier. Such a behaviour is expected, as we know that endocytosis cannot occur in dead cells, and some active element is necessary to induce entry.

### The Importance of Co-receptors

In FIG.3, we also investigate the importance of coreceptors. Co-receptors are a key part of endocytosis, playing an important role in intracellular signalling[26]. However, our research suggests they play not only an important signalling role, but also a thermodynamic role, with the increase in attractive energy that is induced upon binding to a co-receptor being critical to overcome the energetic barriers required for cell entry.

In FIG.3a, we can see that the presence of co-receptors massively decreases this energetic barrier. For example, with 3000 chains *μ*m^−2^, and a spontaneous curvature fraction of 1, HIV-1 has an energy barrier of around 50 kT in the absence of CCR5/CXCR4, which drops to almost zero in the presence of co-receptors. Likewise, the presence of co-receptors massively lowers the HS density required for entry by orders of magnitude, as indicated in FIG.3b.

### Viral Load and Entry Probability

Using this energy barrier we can estimate the probability that a bound particle will enter the cell. We can define this probability using the Boltzmann factor, as

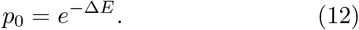

We can then express the probability that any virus will enter the cell in terms of the number of virions bound to the cell surface, *n_V_* as

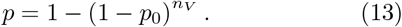

The number of viruses bound to the cell can be determined using the surface coverage *θ*, as

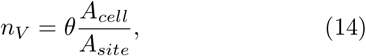

where *A_site_* is the area of a binding site, and *A_cell_* is the surface area. For the epithelial cells attacked by SARS-CoV-2, this is around 1000 *μ*m^2^; while for the helper T cells attacked by HIV-1, this is around 300 *μ*m^2^. The surface coverage can be calculated using the energy of the system prior to wrapping, *E*_0_ = *E*(*h* = 0), and the binding contributions between the spike and HS chains[4], as

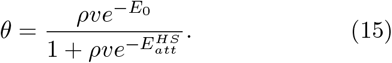

Here *v* is the binding volume, defined as 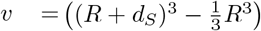, and *ρ* is the viral load, i.e., the number of viral particles per unit volume (usually given in units of copies per mL). The binding energy between a single SARS-CoV-2 spike and HS, *ΔG_HS_* is around −23 kT[4], while for HIV-1 the binding energy is around −6 kT [14].

Using the model outlined above, we then investigated how the probability of entry varies as a function of viral load FIG.4a. For HIV-1, we see an almost linear relationship between viral load and entry probability until becoming asymptotic around 1, while SARS-CoV-2 has an entry of 1 for all viral loads investigated. This model also reveals SARS-CoV-2 has an almost significantly higher probability of entering a target cell than HIV-1 for lower viral loads. The model agrees with what is known about the transmissibilities of each disease: COVID-19 is very highly transmissible, while HIV has low transmissibility, with a risk per exposure below 1.5% for all transmission methods[27].

**FIG. 4.**
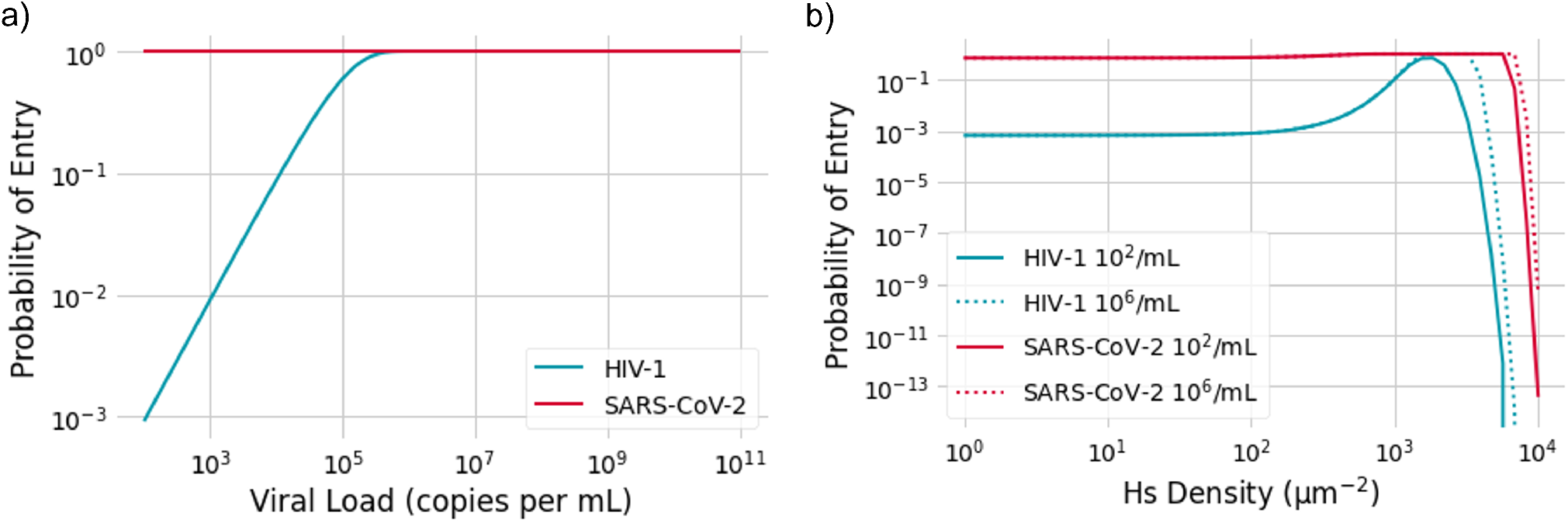
**a)** Viral entry probability as a function of viral load. SARS-CoV-2 has a probability of unity for all viral loads while we see for low viral loads of HIV-1, we have low probabilities of entry, which grow linearly until approaching 1. **b)** Viral entry probability against HS density for low and high viral loads of HIV-1 and SARS-CoV-2. We see for low densities of HS, entry probability is unaffected. As we reach intermediate densities of HS, the entry probability starts to grow due to lowering the potential energy barrier. Then at high densities of HS, the probability falls rapidly to zero, as virions can no longer penetrate the HS brush, and cannot reach the cell surface.

Finally, we plotted the probability of entry for low (10^2^), high (10^6^) viral loads of HIV-1 and SARS-CoV-2 against HS density. We see that the probability of entry for SARS-CoV-2 is unity for all low and medium HS densities and then rapidly approaches zero for high densities.

HIV-1 shows more interesting behaviour, remaining constant for low densities, before increasing in an intermediate range, before rapidly approaching zero for high densities. We can see that for a certain density of HS, between 10^3^ and 10^4^ *μ*m, we see an entry probability of 1.

This behaviour can be explained by consideration of the number of bound viruses. For high densities of HS, the energy required to penetrate the chains and reach the surface will be so high as to make the binding energetically unfavourable, leading to no viruses being present on the surface. However, in the intermediate range, viruses can still accumulate on the surface, and the increasing density of HS chains will decrease the energetic barrier, making viral entry more probable. In our previous work[4], we found that intermediate densities of HS promotes the attachment of viruses, while high densities inhibit the attachment. The above results show that a similar story plays out for endocytosis.

It is worth noting that in this model, we neglect any binding interaction between the virions and the HS chains in the wrapping stage. In other work[4], we have shown that this binding interaction may allow virions to overcome the repulsion of dense HS chains and bind to cells with high HS densities. However, it is unclear how this interaction will affect wrapping, whether it will facilitate or hinder wrapping. In conclusion, we have shown that the glycocalyx plays a key role in determining whether viral entry is possible, and we have further shown that co-receptors play an important thermodynamic role in viral entry. We have created a simple thermodynamic model, which can be used to predict the outcome and severity of viral infections.

## Notes

### Competing Interest Statement

The authors have declared no competing interest.

